# Shoot phenotyping of cytokinin receptors mutants revealed fluorescence parameters as early markers of drought stress

**DOI:** 10.1101/2023.11.30.569457

**Authors:** Ján Šmeringai, Jiří Rudolf, Martin Trtílek, Petra Procházková Schrumpfová, Markéta Pernisová

## Abstract

Plant phenotyping represents an increasing promise in plant research by providing a complex picture of plant development and fitness. In research focused on various environmental stresses, phenotyping can uncover markers that can sensitively assess the stress impact in very early stages before morphological changes. PlantScreen^TM^ System represents a tool dedicated for shoot and root phenotyping in soil enabling high-precision, high-throughput phenotyping of small, mid-size and large plants. The system offers wide range of sensors providing the number of non-invasive analyses of morphological and physiological parameters as well as of pigments, water, or metabolite content.

In our work, we combined phenotyping approaches to determine morphological changes and the status of the photosynthetic apparatus in Arabidopsis plants exposed to drought stress. Focused on morphology, the rosette area became smaller after seven days of drought stress when compared to control conditions. Interestingly, cytokinin signalling mutant *ahk2 ahk3* revealed drought resistance compared to other genotypes. The fluorescent parameters showed higher sensitivity even in wild type. Non-photochemical quenching displayed values connected to reduced activity of photosynthetic apparatus after five days of drought stress. Taken together, acquired fluorescence parameters can serve as a marker of drought stress detection before morphological alterations occur.

**Highlight:** Fluorescence parameters can serve as early markers of drought stress before morphological alterations appear. Shoot phenotyping of cytokinin receptor mutants showed drought resistance in the *ahk2 ahk3* double mutant.

## Introduction

The green revolution in the first half of the 20^th^ century, helped increase world food security sharply, which allowed higher standards of living, and a rapid increase in the global population. These accomplishments, in their turn, gave rise to many new challenges. Intensive agriculture has resulted in soil exhaustion, soil degradation and, decrease in biodiversity (Tsiafouli *et al*., 2015). Alongside that, the climate change is increasing the frequency of extreme weather phenomena such as heatwaves, droughts or heavy precipitation (Rummukainen, 2012). Plants, being sessile organisms, are inevitably exposed to these weather fluctuations, which can result in the disruption of their physiology and morphology. Since food production is vastly dependent on proper functioning of the plant body, these challenges pose a major barrier for sufficient food production. Therefore, while basic research should aim at uncovering the defence mechanisms used by plants to withstand stress conditions (Hu and Xiong, 2014), applied research should urgently focus on breeding (and introduction into agricultural practice) of more crop varieties resistant to biotic or abiotic stresses.

Non-destructive and repeatable methods that could be used on large-scale experiments is absolutely essential to achieve these stated goals. Recent technical advances have provided us with a broad set of tools used for plant body phenotyping, which is defined as the set of methodologies and protocols used to precisely measure plant growth, architecture, and composition at different scales, in controlled environments and in the field (Fiorani and Schurr, 2013; Yang *et al*., 2014; Milella *et al*., 2019; Mertens *et al*., 2021). The necessity to perform these measurements in large numbers, means that high-throughput phenotyping is now seen as a major bottleneck in the association of crop improvement and information regarding phenotype data (Feng *et al*., 2017). This problem was partially resolved with several integrating technologies over the past two decades. However, the high dimensional datasets provided by phenotyping methods are a challenge for data processing and evaluation, which has become a new bottleneck in the whole process (Minervini *et al*., 2016). Deep learning has been proposed as a solution to alleviate this issue (reviewed in (Arya *et al*., 2022)).

Measuring the kinetics of chlorophyll *a* fluorescence is a useful non-invasive method for evaluating the condition of the photosynthetic apparatus, and is often used in phenotyping (Sperdouli *et al*., 2021; Shomali *et al*., 2023). Energy absorbed by photosystem II can be utilized in three main ways: i) it can be used up in photosynthetic processes (photosynthetic quenching); ii) dissipated as heat (non-photochemical quenching) or iii) re-emitted as fluorescence back into the environment (reviewed in (Roháček and Barták, 1999). Rapid light curve is a method of measuring induced fluorescence kinetics often used for assessing stress in plants in controlled environments or in the field (Rascher *et al*., 2000; Kalaji *et al*., 2016; Sánchez-Reinoso *et al*., 2019). To measure the rapid light curve, the plants are exposed to gradually increasing intensities of light. This allows us to obtain useful parameters such as maximum quantum yield (F_V_/F_M_), actual quantum yield of photosystem II (QY) or non-photochemical quenching (NPQ). RGB image analysis is used for phenotyping the plants. This method can be based on the analysis of the colour of individual pixels to provide us with useful information about the morphology of the observed plants (Pavicic *et al*., 2017). Simultaneously, the data can also be used for the analysis of the presence and composition of individual pigments in the plant shoot (Bednaříková *et al*., 2023). Higher-resolution and more complex information can be obtained by combining pigment analysis, chlorophyll fluorescence and morphology. The PlantScreen^TM^ System is capable of high-throughput, high-precision, and non-invasive measurement of the plant phenotype, combining RGB analysis of shoot morphology and colour spectrum, measurement of chlorophyll fluorescence kinetics and image analysis of the root system. Such a combination of phenotyping techniques provides comprehensive information about the plant – comprising information on growth and development, physiological status and fitness, yield and biomass production or reactions to stresses.

Biotic and abiotic stress are some of the main causes for decrease in agronomical yield (Pandey *et al*., 2017; Kopecká *et al*., 2023) and therefore, breeding has focused on the development of new approaches to improve crop stress tolerance (González Guzmán *et al*., 2022). These approaches require a robust understanding of plant physiology in terms of stress responses and especially plant hormone signalling, which controls plant development and ultimately, crop yield (Sah *et al*., 2016; Wang *et al*., 2021; Castro-Camba *et al*., 2022; Mandal *et al*., 2022*a*).

Cytokinins play an important role in stress pre-adaptations and stress tolerance (Mandal *et al*., 2022*a*,*b*). Modulation of cytokinin biosynthesis leads to increased drought tolerance (Reguera *et al*., 2013). Furthermore, cytokinins are crucial regulators of hydrotropism in several species, which suggests even tighter link between drought stress resistance and cytokinin signalling (Chang *et al*., 2019; Miyazawa and Takahashi, 2020).

Cytokinin signalling employs the multistep phosphorelay which helps the plant to integrate various environmental stimuli, including abiotic stresses (Skalak *et al*., 2021). In Arabidopsis, the multistep phosphorelay comprises three families of proteins – histidine kinases (AHKs), His-containing phosphotransfer proteins (AHPs) and response regulators (ARRs), which can be functionally and structurally sub-divided into three groups (A-ARRs, B-ARRs, C-ARRs) (reviewed in (Kieber and Schaller, 2018). The AHKs represent a diverse group of proteins which can act in both a cytokinin-dependent or independent manner depending on differences in their domain composition (Spíchal *et al*., 2004; Tran *et al*., 2007; Desikan *et al*., 2008; Deng *et al*., 2010). The cytokinin receptors AHK2-4 (Inoue *et al*., 2001; Higuchi *et al*., 2004; Nishimura *et al*., 2004) possess the cytokinin-binding CHASE domain (Anantharaman and Aravind, 2001) followed by a transmembrane domain, a histidine kinase domain and a receiver domain. Upon activation, AHKs autophosphorylate on their histidine kinase domain and trigger the His-Asp phosphotranspher toward the conserved Asp residue of their receiver domains. The phosphate group is further transferred to AHPs and finally to B-ARRs – transcription factors , which are activable by phosphorylation (Sakai *et al*., 2000; Argyros *et al*., 2008) and A-ARRs, which act as major negative feedback links in the signalling pathway (To *et al*., 2004; Kim, 2008). The precise functioning of C-ARRs has not been elucidated yet. They presumably act as AHP-phosphatases with a positive impact on abiotic stress tolerance, as it was shown that the expression of ARR22 increases when exposed to drought and freezing (Gattolin *et al*., 2006; Horak *et al*., 2008; Kang *et al*., 2013).

Regarding stressors, cytokinin receptors AHK2, AHK3, and AHK4 negatively regulate drought and dehydration tolerance, as seen by the phenotypes of their corresponding mutants (Tran *et al*., 2007; Kang *et al*., 2012). Similarly, AHPs and B-ARRs act as partially redundant negative regulators of drought tolerance (Nishiyama *et al*., 2013; Nguyen *et al*., 2016). Complementarily, the negative components of cytokinin signalling – A-ARRs and C-ARRs have been observed to positively affect water-deficiency stress tolerance (Wohlbach *et al*., 2008; Kang *et al*., 2013; Huang *et al*., 2018). Interestingly, although the cytokinin receptors seem to act as negative regulators of drought tolerance, the cytokinin-independent AHK1 is a positive effector of water-deficiency tolerance and is a potential osmosensor (Tran *et al*., 2007; Wohlbach *et al*., 2008; Kumar *et al*., 2013). Further, AHK5 negatively regulates plant tolerance to osmotic and drought stress, possibly due to its role in ROS-dependent stomatal closure and crosstalk with ethylene and ABA signalling (Iwama *et al*., 2006; Desikan *et al*., 2008; Pham and Desikan, 2012; Pham *et al*., 2012; Szmitkowska *et al*., 2021).

In summary, with respect to drought tolerance, the positive executors of multistep phosphorelay seem to act as negative regulators, while the quenching components seem to act as positive regulators. Although the components are partially redundant, their function does not overlap completely. This presents a conundrum which can potentially be solved by precise phenotyping combined with appropriate data analysis, which is now possible using the PlantScreen^TM^ Compact System. Since the system generates large amounts of data, the main concern is data multidimensionality – with all the associated advantages and disadvantages. The main concerns include data visualization, reduction of dimensionality and further simplification without losing information.

In our work, we implemented and optimised a drought stress protocol on an assembly of Arabidopsis wild type (Col) and single and double mutants in the cytokinin receptors AHK2 and AHK3. Aiming for proper data analysis, several methods of multivariate statistical analysis were employed covering clustering analysis, ordination methods like principal component analysis (PCA), and a machine learning approach (random forest) to enable the extraction of important parameters. Our results indicate that the drought impact can be significantly differentiated from control within 5 days of drought stress using fluorescence parameters. Non-photochemical quenching and qN were identified as the most important parameters and indicated the involvement of protective mechanisms of the photosynthetic apparatus during drought stress. Comparing wild type and cytokinin receptor mutants, parameters from all three inspected categories (morphology, color segmentation and chlorophyll fluorescence) showed a high degree of importance. Altogether, combining multiple phenotyping approaches and advanced data analysis furnishes a comprehensive set of information about plant morphology, physiology, and fitness during drought stress. Fluorescence parameters were shown to be useful markers of the very early phases of drought stress before morphological changes are apparent. This protocol can now be applied to study other stresses, cultivation conditions or genotypes.

## Materials and methods

### Plant material

All plant material used was *Arabidopsis thaliana*. The Nottingham Arabidopsis Stock Centre (NASC) provided seeds for the wild type accession Col (N60000). The mutant lines have been described previously: *ahk2-5*, *ahk3-7*, *ahk2-5 ahk3-7* (Riefler *et al*., 2006). The *AGI* codes (www.arabidopsis.org) are: *AHK2* (AT5G35750), *AHK3* (AT1G27320).

### Growth conditions

For plant cultivation we used the Klasmann Substrate TS3 fine (416, Pasič) and 7×7×6.5 cm pots (Pasič). Seeds were geminated in one pot and were replanted after 7 days in the very centre of a single pot. Plants were grown in cultivation chambers – phytotrons (CLF Plant Climatics or PSI - Photon Systems Instruments), under long-day conditions (16 hours light/8 hours dark) at 21°C with a light intensity of 150 μMm^−2^ s^−1^ and 40-60% relative humidity. Watering was adjusted according to control or drought stress conditions (Fig. 1A).

**Fig. 1.**
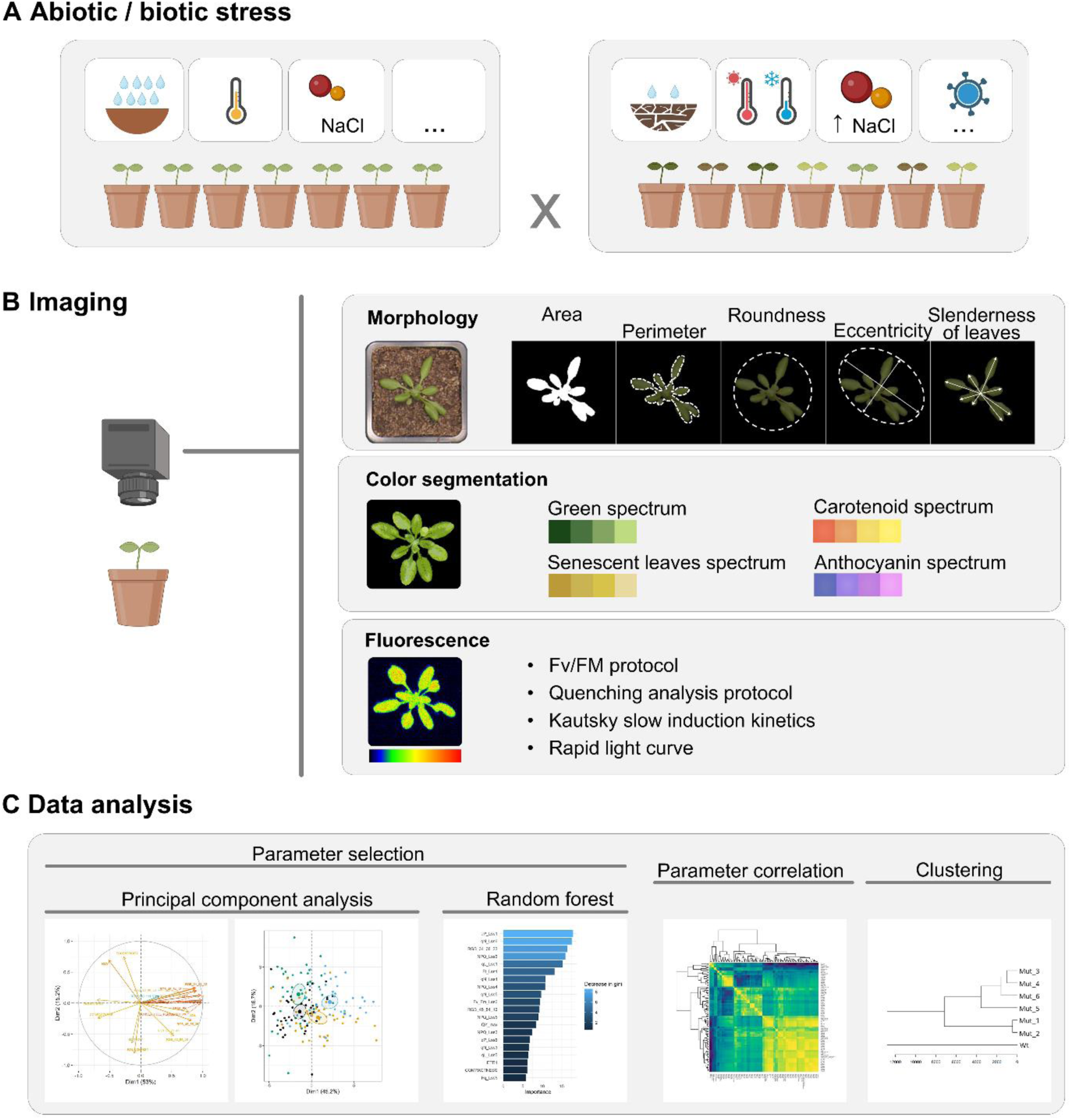
Drought stress experimental setup and phenotyping using PlantScreen^TM^ Compact System. (A) Potential abiotic or biotic stress is applied to plants. (B) Imaging by PSI DUAL CAM or by side RGB cameras provides raw data used for calculating phenotype parameters. RGB cameras enable extraction of morphological parameters (e.g., area, perimeter, roundness, eccentricity, slenderness of leaves) or colour segmentation. Colour segmentation allows the definition of any colour from the RGB spectrum and can detect senescent leaves or anthocyanins. Fluorescence camera allows imaging of the chlorophyll fluorescence signal with various measuring protocols including F_V_/F_M_, quenching analysis, Kautsky slow induction kinetics or rapid light curve. (C) Downstream analysis of the parameters based on multidimensional principal component analysis or machine learning approach like random forest is used to estimate the importance of morphological, colour or fluorescence parameters. Correlograms utilizing Spearman rank correlation visualize correlating parameters. Further clustering of parameters, such as genotypes, treatments, or stress conditions, helps identify the impact of the stresses on plants. Created with BioRender.com.

### PlantScreen^TM^ Compact System

#### Hardware

The phenotype data were acquired using the PlantScreen^TM^ Compact System (developed by Photon Systems Instruments, Drásov, Czech Republic), which is designed for digital phenotyping and cultivation of small and mid-size plants up to 0.8 m in height (e.g., *Arabidopsis thaliana*, cereals, strawberries, turfgrass, young soybean, tobacco, corn, etc.). In our version the transport of plants is carried out manually in trays that can be adapted to carry different patterns for single or multiple plants grown in individual pots or *in vitro* (e.g., multiwell plates) providing flexibility of use with numerous different species, or with a single species throughout its growth cycle. The entire system is built in a light isolated box with active internal ventilation, which does not transmit any ambient light from outside. For precise morphometric imaging and correct measurement of chlorophyll fluorescence, dark/light adaptation is included. Environmental parameters are monitored using temperature, humidity, and light sensors.

The system is equipped with several imaging sensors:

i. Dual (monochromatic and RGB) camera for top view RGB and chlorophyll fluorescence imaging – the PSI DUAL CAM is fitted with two 12.36-megapixel CMOS sensors, a monochromatic Sony IMX253LLR-C for chlorophyll fluorescence measurement and a colour Sony IMX253LQR-C sensor for RGB structural imaging. These sensors deliver a resolution of 4112 × 3006 pixels as well as global shutter feature. The sensors are extremely sensitive and are real megapixel CCD substitutes that produce sharp and low-noise images. The monochromatic sensor runs in binning mode with 2056 x 1503 resolution, which means four times higher sensitivity.
ii. RGB camera for side view imaging (linear scan) – the UI-5580CP is fitted with a 5-megapixel CMOS sensor. About half an inch in size, the sensor delivers a resolution of 2560 x 1920 pixels as well as rolling and global start shutter features. The various shutter modes produce sharp, low-noise images.
iii. RGB camera for root imaging – the PSI BW is fitted with a 12.36-megapixel CMOS sensor (Sony IMX253LQR-M). The sensor delivers a resolution of 4112 × 3006 pixels as well as global shutter feature.

#### Software

The comprehensive software package with remote accessibility comprises the PlantScreen™ Server application, the PlantScreen™ Scheduler client, a PlantScreen™ Database and a PlantScreen™ Data Analyzer. The package provides control over all imaging modules, as well as database configuration, data acquisition and image analysis. A set of computed morphological parameters is available: area, perimeter, compactness, rotational mass symmetry (RMS), roundness, isotropy, eccentricity, slenderness of leaves (SOL) and convex hull area (Fig. 1B). A detailed definition of the morphological parameters has been published previously (Pavicic *et al*., 2017).

Several protocols for fluorescence measurement are available:

i. F_V_/F_M_ protocol – protocol providing two measured parameters F_0_ and F_M_, and one calculated parameter, maximum QY. Detailed definition of parameters can be found in supplementary data (Supplementary Table S1).
ii. Kautsky slow induction kinetics – during this protocol, plants in the dark-adapted state are exposed to actinic irradiation, which results in a rapid increase in fluorescence culminating in maximum F_P_ value. With ongoing exposition to actinic light, the photosynthetic apparatus accommodates to the radiation via the involvement of secondary photosynthetic processes which results in reduced fluorescence until transient fluorescence (F_T_) is reached (Roháček and Barták, 1999). Out of the calculated parameters F_V_, QY_max and Rfd are available. Detailed definition of parameters can be found in supplementary data (Supplementary Table S2).
iii. Quenching analysis protocol – Kautsky kinetics supplemented with saturation pulses. The plants in the dark-adapted state are firstly exposed to a saturation pulse and afterwards, actinic light is switched on. Saturation pulses are then applied on plants in the light-adapted state throughout the measurement. This protocol provides a broad spectrum of measured (F_0_, F_M_, F_P_, F_M__Ln, F_M__LSS, F_T__Ln, F_T__Lss) and calculated (F_V_, F_0__Ln, F_0__Lss, F_v__Ln, F_v__Lss, F_q__Ln, F_q__Lss, QY_max, F_V_/F_M__Lss, F_V_/F_M__Ln, QY_Ln, QY_Lss, NPQ_Ln, NPQ_Lss, qN_Ln, qN_Lss, qP_Ln, qP_Lss, qL_Ln, qL_Lss, Rfd_Ln, Rfd_Lss) parameters. Detailed definition of parameters can be found in supplementary data (Supplementary Table S3).
iv. Rapid light curve – plants in the dark-adapted state are exposed to a saturation pulse and subsequently exposed to low intensity actinic irradiation. After a certain time, another saturation pulse is applied on plants in the light-adapted state. This saturation pulse is followed by exposure to actinic light whose intensity has been increased by a specific amount (White and Critchley, 1999). This process and the increase of radiation intensity is repeated six times. This analysis provides a broad range of measured (F_0_, F_M_, F_P_, F_T_) and calculated (F_V_, F_M__Lss, F_T__Lss, F_0__Lss, F_v__Lss, F_q__Lss, QY_max, F_V_/F_M__Lss, QY_Lss, NPQ_Lss, qN_Lss, qP_Lss, qL_Lss, ETR_Lss) parameters. Detailed definition of parameters can be found in supplementary data (Supplementary Table S4).

### Acquiring RGB parameters

The plants in pots were arranged in transportable trays, each of which held 20 plants (4x5 template). The top RGB camera was used to acquire the morphology parameters. The obtained images were pre-processed via the PlantScreen^TM^ Data Analyzer software to get RGB and binary data. These data were used to calculate the morphological parameters (Fig. 1B).

The RGB data were used for colour segmentation analysis. Each pixel was indexed assigned to the colour map according to its R, G, B channel ratio. These values were categorised into groups according to the hues (Awlia *et al*., 2021). The R, G, B values were grouped into 9 hues (R110, G111, B90; R90, G98, B58; R72, G84, B58; R73, G86, B36; R57, G71, B46; R59, G71, B20; R45, G55, B36; R45, G54, B13; R34, G38, B22) (Fig. 1B).

### Acquiring fluorescence parameters

The physiological status of plant lines was assessed by measuring the induced fluorescence of chlorophyl *a* (Fig. 1B). We used Rapid Light Curve protocol, which is suitable for detecting stress affecting plants (Flexas *et al*., 1999). The measurement was performed at a working distance of 430 mm on plants in the dark-adapted state. Arabidopsis plants were kept in the dark for 7 minutes. Firstly, the measuring light was applied for 5 seconds to acquire information about the minimum level of fluorescence in the dark-adapted state (F_0_). A saturation pulse was applied for 800 ms, its source is a LED, which has cool white 6500K spectrum, corresponding almost exactly to daylight, and the intensity of the saturation pulse was 1256 µmol.m^-2^.s^-1^. This pulse with high irradiation intensity is used to determine the maximum fluorescence in the dark-adapted state (F_M_). The actinic light intensities of the individual steps were 15.6, 193.4, 396, 602, 810, and 1013 µmol.m^-2^.s^-1^. Each step begins with a 800 ms saturation pulse in order to measure maximum fluorescence in the light-adapted state (F_M__Lss). Subsequently, the actinic light is switched on for 60 seconds, and at the end of each step the transient fluorescence (F_T_) is measured. The acquired data of the measured fluorescence were transferred to the PlantScreen^TM^ Data Analyzer, which provided the calculated fluorescence parameters.

### Data analysis and statistical analysis (Fig. 1C)

#### Correlograms

The graphical representation of variable correlations was used to detect clusters of partial redundant variables present in the dataset since the variable correlation is one of the prerequisities for efficient principal component analysis (Supplementary Fig. S1). In order to achieve a more robust representation, Spearman correlation coefficient was used, and the representation was colour-coded using the viridis colour palette. Further, the variables were hierarchically clustered based on their correlations using average agglomeration. The correlograms were made using the *stats* package (R Core Team, 2023).

#### Principal component analysis (PCA)

PCA as one of the ordination methods was used to visualise the data and simplify its multivariate nature. The data were scaled and subset into fluorescence-based and RGB-imaging-based parameters, the latter consisting of morphometric parameters and colour segmentation variables. The number of principal components was set based on a scree plot evaluation (Supplementary Fig. S2A, B). The quality of the representation was evaluated by cos^2^ (Supplementary Fig. S2C, D). For the sake of clarity, biplot was used as the final representation only in the demonstrative case. The package *factoextra* (Kassambara and Mundt, 2020) was used. For visualisation of differences between defined groups (genotype vs. stress conditions), the centroids with ellipses representing their 95% confidence interval were depicted in the plots.

#### Random forest

A random forest algorithm was used for estimating parameter importance, utilising *randomForest* package (Breiman, 2001; Liaw and Wiener, 2002). The models were trained on the subset data covering all measured and derived variables. The number of trees was set to 1000, each split tested 9 variables (based on square root estimation). The variable importance was extracted to enable prediction of stress conditions or genotype (i.e., classification model). The performance of the model was tested by confusion matrices using randomly subset test data, which were not used for model training (Supplementary Tables S5-S8). The 20 most important variables were visualised by *vip: Variable Importance Plots* package (Greenwell and Boehmke, 2020) with estimated importance on the main axis and colour coded mean decrease in gini.

#### Hierarchical clustering

In order to see similarities in the tested genotypes under stress conditions, the averaged values for each group were clustered based on Euclidean distance.

#### Time series plots

The selected parameters are depicted as a time series for each genotype under both tested conditions. The lines represent LOESS regressions with shaded 95% confidence interval. The images were generated by *ggplot2* and *ggpubr* (Wickham, 2016; Kassambara, 2022).

#### Time point plots

The selected parameters at representative time points are depicted by Tukey’s boxplots. The images were generated by *ggplot2* and *ggpubr*. The statistical significances were evaluated with a Krustal-Wallis test and a post-hoc Dunn test. The significant differences were depicted using compact letter display. The packages used for this were *rcompanion* (Mangiafico, 2023), *FSA* (Ogle *et al*., 2023), and *stats* (R Core Team, 2023).

Data management, storage and access to users was managed in collaboration with the Biological Data Management and Analysis Core Facility (Svoboda *et al*., 2023).

## Results and discussion

### Optimization of the drought stress experimental setup

It is necessary to ensure precise regulation of water content in the substrate to determine the effect of drought stress on the morphological and physiological aspects of plants. Cultivation conditions were optimized to set the appropriate substrate humidity suitable for the drought stress experimental protocol. The substrate was aliquoted at 50 g per each pot. The commercially available substrate already had a humidity of 54%. After preparing the pots, we added 15 ml of water (10 ml before replanting the seedlings and 5 ml afterwards) to keep the transplanted roots wet. Thus, the overall humidity of the substrate reached approximately 65%. 11 days after sowing (DAS), the first phenotyping was performed to verify uniformity of the observed parameters in plant lines before applying drought stress. At this timepoint, both variants were equal in substrate water content. After the first phenotyping, no more water was added to the substrate for the drought stress variant. The phenotyping was then carried out three times per week at 14, 16, 18, 21, 23 and 25 DAS corresponding to 3, 5, 7, 10, 12, 14 days of drought stress (DODS; Fig. 2A).

**Fig. 2.**
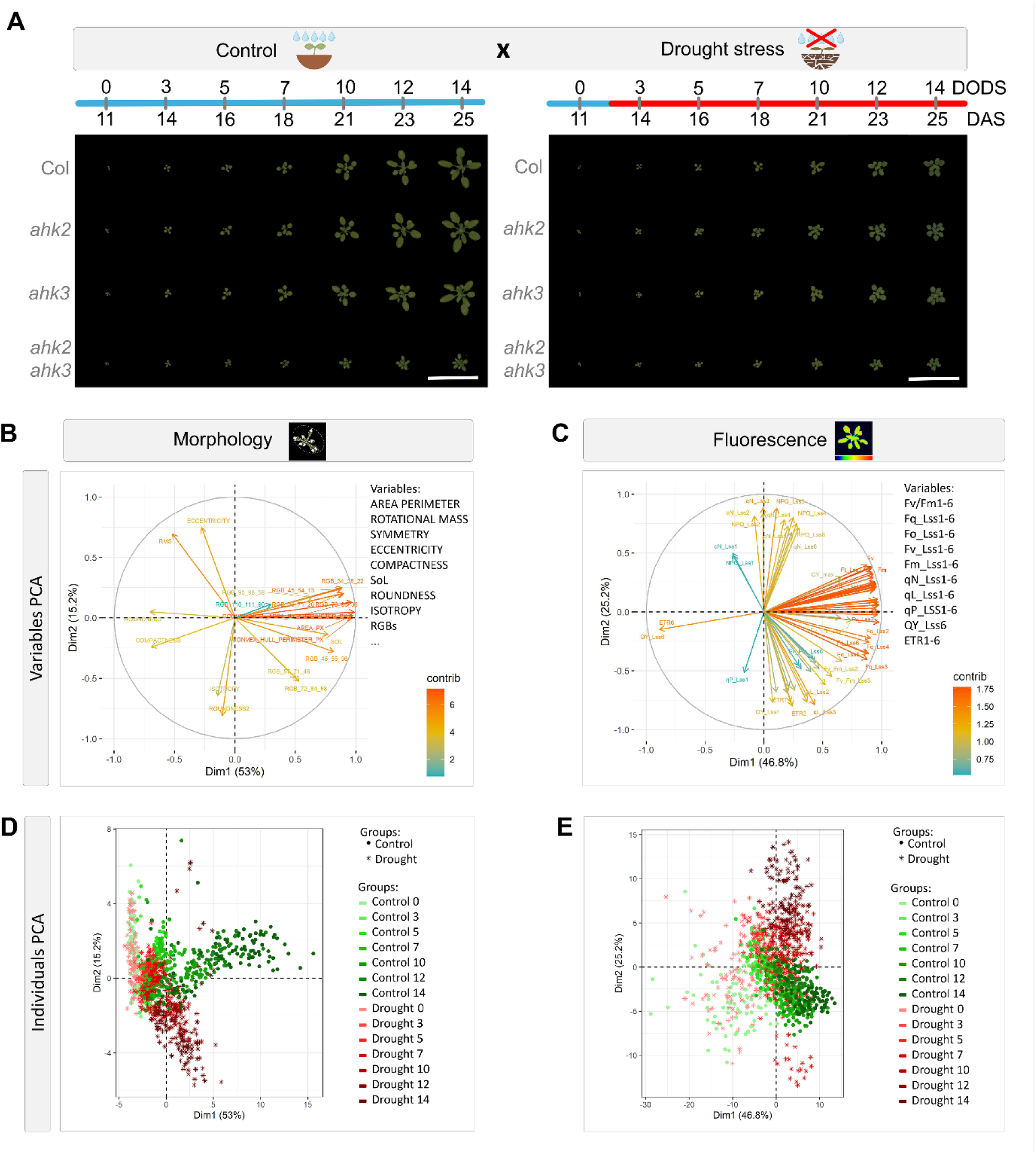
Measurement of morphological and fluorescence parameters in wild type (Col) and cytokinin receptor mutants (*ahk2*, *ahk3*, *ahk2 ahk3*). (A) Plants were exposed to drought stress and phenotyped for 14 days (0-14 DODS). The age of plants corresponds to days after sowing (DAS). Scale bar: 7 cm. (B, C) Charts of variable contribution to PCA models were built using (B) RGB-based parameters or (C) fluorescence imaging. Variable labels are shown in simplified form. Detailed version is available as supplementary data (Supplementary Figure S3). (D, E) In the corresponding space, single data points are depicted showing global differences with progressing drought stress, generated from (D) RGB or (E) fluorescence data.

The effect of severe drought stress on overall plant body fitness has been studied intensively, but the effect of mild and moderate drought stress is far less understood. In our optimized protocol we concentrated on mild stress and the experiment was terminated when severe stress responses began to appear. Mild and severe drought stress seem to activate different mechanisms. In drought resistant barley exposed to mild drought stress, stomatal conductance was the main factor that limited the rate of photosynthetic processes, while in case of severe drought light-dependent reactions were structurally and biochemically impaired (Ghotbi-Ravandi *et al*., 2014). Such processes limit the ability of plant to produce biomass and result in yield reduction (Torres and Henry, 2018). Moreover, it seems that there are no efficient mechanisms to resist moderate drought stress in Arabidopsis, as revealed by the measurement of induced chlorophyll fluorescence, which showed a higher impact on photosynthesis in mild and severe stress when compared to moderate stress. The mild stress reaction was comparable to the severe stress effect (Sperdouli and Moustakas, 2012).

### The data set involves multiple collinear variables and shows global differences in stressed plants

Our results revealed several parameters to be highly correlated (Supplementary Fig. S1). This is especially valid for parameters acquired at a range of different light intensities (different LSSs). Moreover, this multicollinearity can be explained by the fact that the dataset involves both directly measured and calculated parameters and therefore a certain level of correlation can be expected between the derived ones and their corresponding source.

Since PCA can effectively simplify the data with collinearity (Lafi and Kaneene, 1992), this ordination method was selected for global data visualisation (Fig. 2B-E). Two models were created on a dataset comprising all data points, where variables were divided into two groups based on their acquisition origin. The first model was created on RGB-based variables, which involve morphometric parameters and colour segmentation (Fig. 2B). The second model was created using fluorescence-based parameters (Fig. 2C). Two principal components were selected for both models, which in both cases cumulatively explained approximately 70% of the original variance. The contribution of each parameter was evaluated (Fig. 2B,C).

Further, individual data points which represent measurements from a single plant at each time point are depicted in the selected 2-component plane (Fig. 2D,E). The data points are colour-coded with a light colour representing the experiment initiation and a progressively darker colour corresponds to progressing time. The non-stressed control is depicted in shades of green/points, whereas samples stressed by drought are depicted in shades of red/asterisks. The non-stressed and stressed clusters differentiated in the later phases of the experiment, which correspond to morphological, colour, and fluorescence changes which occurred in response to drought. In conclusion, it is possible to distinguish the global effects of drought stress on the plants. However, this methodology does not provide information about variable importance, which was further estimated applying a random forest approach.

### Fluorescence parameters as markers of drought stress

The random forest is a powerful machine learning algorithm which can be used for classifications, regressions, and measurements of variable importance (Genuer *et al*., 2010). We used a random forest algorithm for the classification of drought stress conditions and subsequent estimation of variable importance. The created models were used for parameter importance evaluation, and not as predictors, thus the workflow was adjusted accordingly. The data set comprising all time points was used as a training set for the classification model, which aimed to differentiate between control and stressed plants across the tested lines (Col, *ahk2, ahk3,* and *ahk2 ahk3*). Error rates of the model were 6.23% and 7.41% on the trained dataset and on the test dataset respectively (Supplementary Table S5). This result shows that plants exposed to drought stress can be differentiated from non-stressed controls.

We focused on the earliest time point where stressed plants were clearly differentiable from controls. The model was therefore trained on a subset of data divided based on days since the start of the stress treatment (0, 3, 5, 7, 10, 12, 14 DODS). Most importantly, the classification error was less than 5% for plants after 5 DODS (2.27% train data, 0% test data; Supplementary Table S6) and it further dropped to 0% at 14 DODS (0% train data, 0% test data; Supplementary Table S7), representing the last measuring day. This hints that the impact of drought stress is likely clearly visible as early as 5 days with a reasonably acceptable classification error which then further drops as drought-stressed plants continue to diverge from the watered control. This is consistent with the PCAs (Fig. 2D, E) showing a clear separability of stressed plants from the control at later timepoints. Based on the model, the fluorescence parameters were selected as the most important parameters for stress distinction at 5 DODS. Interestingly, in the late phase of the experiment (14 DODS), the fluorescence parameters were still evaluated by the model as the most important. This was very probably caused by the *ahk2 ahk3* morphology *per se* and therefore the morphological parameters are probably not generally applicable for such analyses. However, fluorescence parameters seem to be appropriate even for this type of experiment and can serve as markers of drought stress.

The effect of drought stress on the physiological status of plants was evaluated by measuring induced chlorophyll fluorescence. The random forest analysis showed that parameters associated with the protective mechanisms of the photosynthetic apparatus [mainly NPQ and qN (Fig. 3A, B)] were among the most important factors that explained the drought effect in plants both at 5 DODS and 14 DODS. NPQ is steady-state non-photochemical quenching, whereas qN is the coefficient of non-photochemical quenching. Both, qN and NPQ point at the quenching mechanisms that play an important role in the estimation of the absorbed energy that is consumed by protective mechanisms, mainly through heat dissipation. NPQ started to increase after exposure to drought stress sooner (5 DODS; Fig. 3C) than qN (8 DODS; Fig. 3D). At both 5 and 14 DODS, qN measured under various light intensities was more important in assessing the drought impact than NPQ (Fig. 3A, B). In higher plants non-photochemical quenching is mainly based on the changing pH level in the thylakoid lumen, which activates the xanthophyll cycle (Horton *et al*., 2000; Müller *et al*., 2001), and is therefore responsible for the de-excitation of chlorophyll molecules in photosystem antennae (Havaux *et al*., 2007). At higher light intensities, NPQ is sensitive to the effect of drought stress (Yao *et al*., 2018), which makes this parameter a useful marker of the early signs of water deficit (Chou *et al*., 2017). With increasing drought, defense mechanisms are activated to protect the photosynthetic machinery and NPQ values increase (Jia *et al*., 2020). However, in case of intense and severe drought stress, the whole system collapses and NPQ decreases (Wang *et al*., 2018). F_V__F_M__Lss5 is also among the parameters that can be used for estimating drought stress at 14 DODS. The F_V__F_M__Lss5 value started to decrease after 8 DODS in stressed plants, whereas in control plants it kept increasing until the end of the experiment (Fig. 3E). F_V__F_M__Lss is defined as the maximum efficiency of photosystem II in the light-adapted state (Oxborough, 2004) and a decrease in this parameter points at the deteriorating condition of the photosynthetic apparatus. This decrease under abiotic stress was previously recorded *in vitro* (Wang *et al*., 2018) and in substrate (Adhikari *et al*., 2019).

**Fig. 3.**
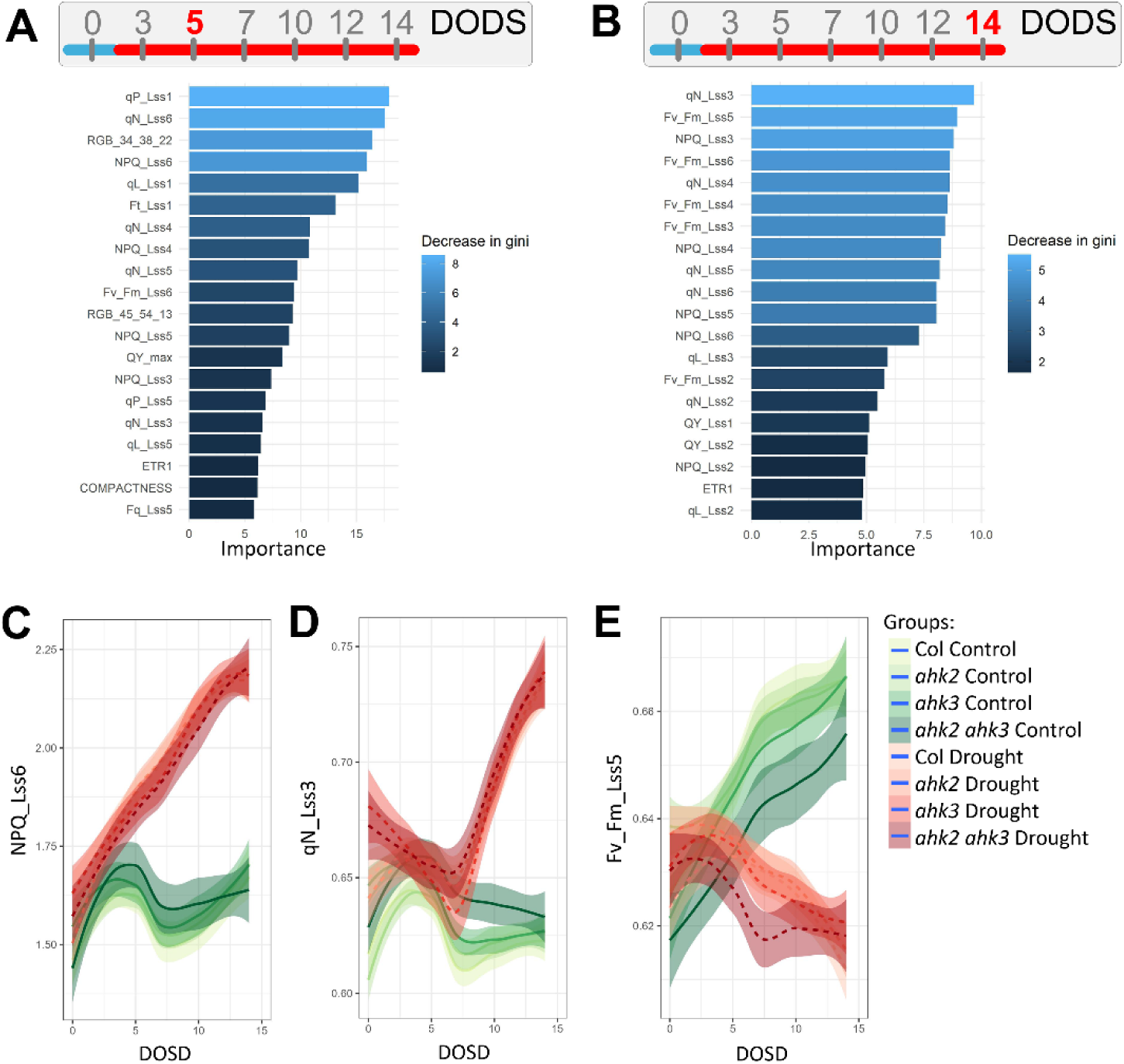
Random forest algorithm was used to identify the most important parameters at (A) 5 DODS and (B) 14 DODS with a color-coded mean decrease in gini. (C-E) Selected important variables are shown as they change during the drought stress in plants exposed to stress (shades of red) and in controls (shades of green). The lines depict LOESS models with a 95% confidence interval in corresponding shades.

### Genotype and cultivation condition clustering reveals distinctive patterns

To look for similarities or differences between the tested genotypes, the dataset was clustered with respect to either genotype or genotype in combination with stress condition (Fig. 4A, B). It was obvious that the biggest difference among the phenotype of the genetic lines is for *ahk2 ahk3*. Col is the most related to the *ahk3* mutant followed by the *ahk2* single mutant. When drought stress is considered, the clustering has a similar fractality. The control and stressed plants belong to different clusters, except for *ahk2 ahk3* which exhibited only small differences between stressed and non-stressed plants (Fig. 4B). In this case, the *ahk2 ahk3* genetic contribution on phenotype is higher than the contribution of drought stress.

**Fig. 4.**
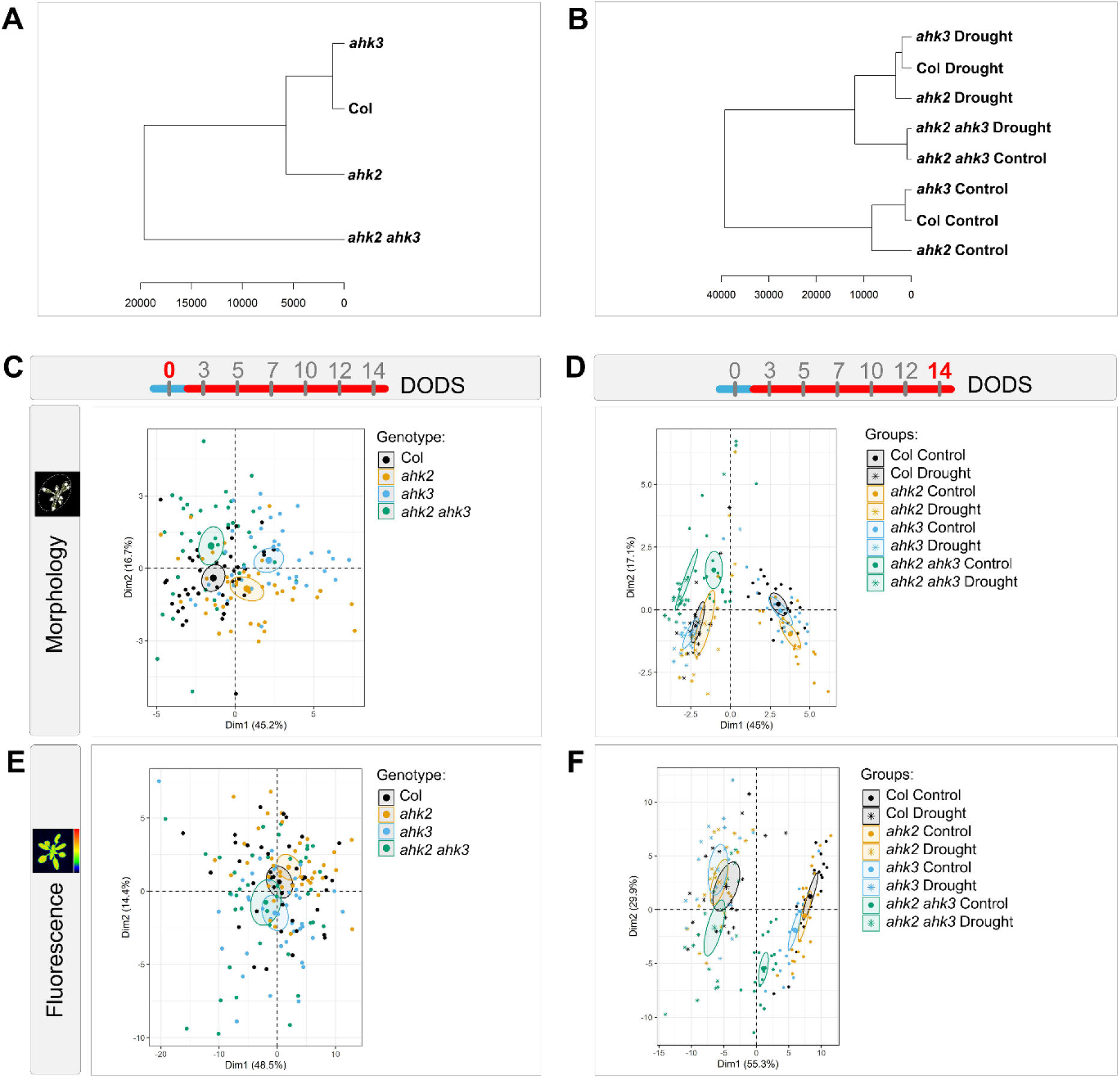
Hierarchical clustering on whole dataset based on (A) genotypes or (B) genotypes in combination with drought stress. (C-F) PCA models were built for (C, E) start of the experiment and (D, F) the end of experiment to detect differences between genotypes at the selected time points. The models were built using (C, D) RGB-based parameters or (E, F) fluorescence-based parameters. All charts show the centroid of each defined group with its 95% confidence interval as a shaded ellipse.

Principal component analysis was performed separately for morphological analysis comprising colour segmentation, and induced chlorophyll fluorescence analysis on the first (11 DAS/ 0 DODS) and the last (23 DAS/ 14 DODS) measurement of the experiment. As for morphology analysis (Fig. 4C), the first analysed phenotype at 11 DAS showed only small morphological differences between the genotypes, since the clusters partially overlap, but no overlaps were detected in the estimations of cluster centroid confidence intervals (depicted as ellipses). This suggests that there are small quantitative differences between the genotypes; nonetheless, the genotypes are not easily separable. Morphological data obtained at 23 DAS revealed a strong distinction between control and drought-stressed plants in the case of Col, *ahk2* and *ahk3* (Fig. 4D). However, no global differences in morphology among Col, *ahk2* and *ahk3* were detected using PCA. A different trend was observed in the case of the *ahk2 ahk3* double mutant, separated by the PCA analysis from other genotypes.

No differences were observed in the PCA analysis of fluorescence parameters at 11 DAS/0 DODS (Fig. 4E), which points to a homogeneity in the rate of primary photosynthetic processes before the drought stress. However, the measurement of induced chlorophyll fluorescence at 23 DAS/14 DODS revealed a strong distinction between well-watered and drought-stressed plants (Fig. 4F). The PCA uncovered a linear separation between control and drought-stressed plants which fits with the result obtained by training a random forest classificator on 14 DODS data showing 0% error rate expected for linearly separable data. Taken together, we were not able to differentiate the individual genotypes exposed to drought stress using PCA. However, in the control group, the cluster of *ahk2 ahk3* mutant was the most distant from other genotypes.

In principle, PCA does not substitute standard statistical methods for the evaluation of differences between genotypes for single parameters, but it points out that the genotypes are very likely not easily separable. This can be further supported by training a random forest classifier, which exhibited approximately 25% error rate (28.91% train data; 25% test data; Supplementary Table S8) in genotype estimation using the data from the last time point measurement. Since the classification error on a random sample should be 75%, the model performed better than randomly generated data. Although not precise, this estimation is better than random and can point out variables which are important for such selection when their biological relevance is carefully evaluated (Fig. 5A).

**Fig. 5.**
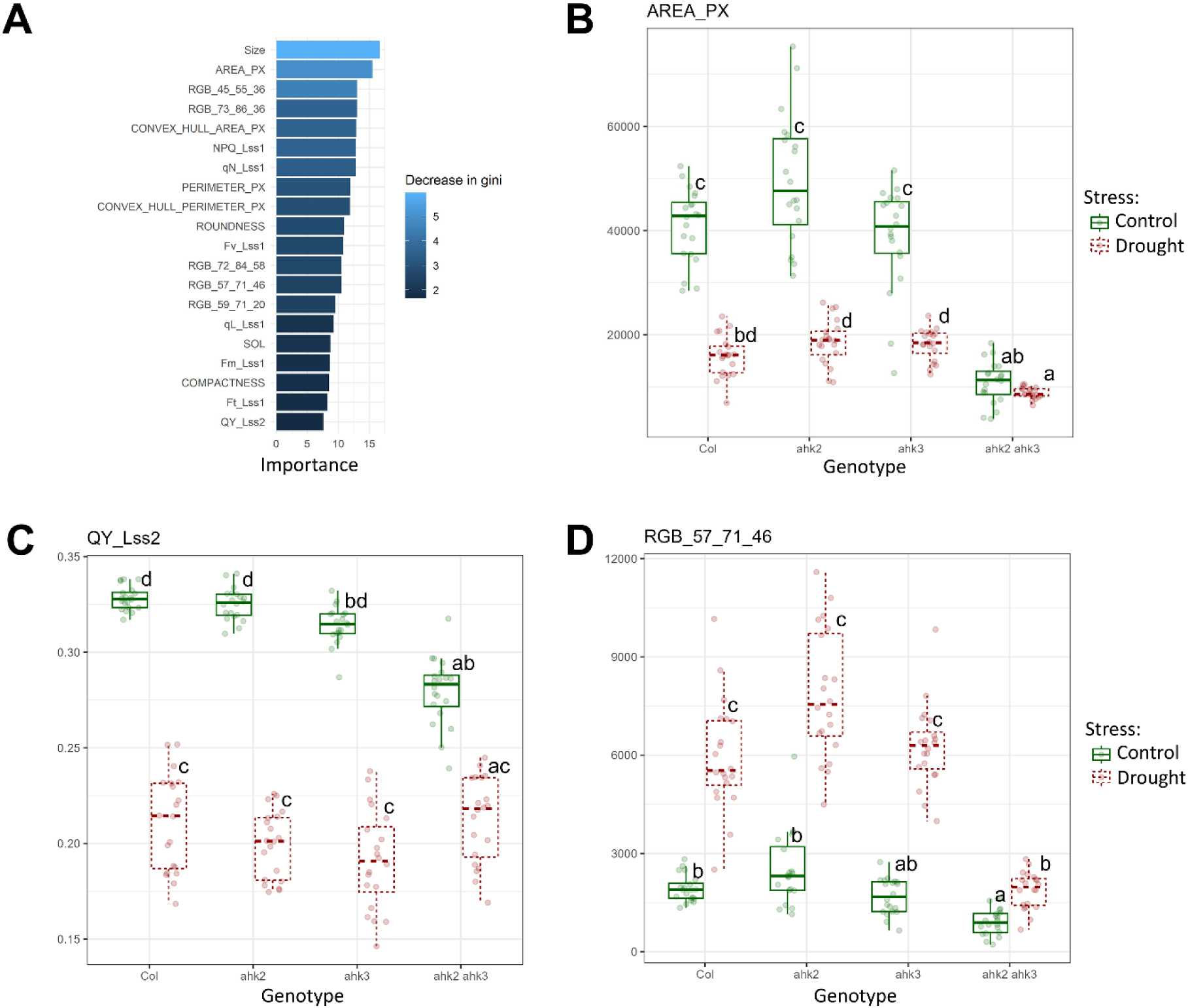
(A) Random forest model was generated to reveal the most important parameters for genotype discrimination using data from 14 DODS. (C-D) Three parameters were selected – (B) one morphological - area, (C) one fluorescence-based - QY, and (D) one colour-segmentation-based – RGB_57_71_46. Lowercase letters in charts indicate significantly different groups as determined using Kruskal-Wallis test and post-hoc Dunn test (p< 0.05).

### The *ahk* mutants exhibit several distinguishable phenotypes

The random forest analysis showed that all categories of parameters (morphological, fluorescence and colour segmentation) showed a high degree of importance for determination of observed genotypes at 14 DODS (Fig. 5A). Apart from parameters suitable for drought stress distinction (Fig. 3), morphological parameters were revealed to be the most important for genotypes distinction, particularly area and size of the plant rosette, calculated as a sum of the pixels showing fluorescence activity during fluorescence measurement. That may be the consequence of the phenotype of the tested plant lines as the *ahk2 ahk3* double mutant displays a much smaller shoot compared to other lines in general (Riefler *et al*., 2006).

Unlike single mutants, *ahk2 ahk3* double mutant is known for its strong phenotype comprising smaller shoot, shorter hypocotyl, compact rosettes with shortened petioles or affected leaf shape (Nishimura *et al*., 2004; Riefler *et al*., 2006). At 14 DODS, no differences in rosette area among wild type plants and single mutants were detected under both control and drought stress conditions (Fig. 5B). In control conditions *ahk2 ahk3* double mutant area is reduced when compared to other observed plant lines. Such a difference in the *ahk2 ahk3* double mutant from other plant lines can be observed also under drought conditions. We also examined the effect of drought on individual genetic lines. Rosette area was reduced when control wild type and single mutants were compared to drought exposed wild type and single mutants. There was no difference between control and drought exposed *ahk2 ahk3* double mutant. Previous research suggests that rosette area can be used as a marker more broadly. It was shown that it is a good determinant of the involvement of mutant lines in the plant body response under various abiotic stresses (Awlia *et al*., 2021).

Chlorophyll fluorescence provides information about the physiological status. Among our results, the most important markers for plant genetic lines variance are NPQ and qN (Fig. 5A), which reflect the proportion of the energy absorbed by the photosynthetic apparatus, that is used up by defense mechanisms. Another important fluorescence parameter with a high degree of importance was quantum yield measured under the effect of a light intensity that was moderately elevated when compared to the cultivation conditions of plants. Quantum yield is one of the most often used fluorescence parameters. Its value describes the amount of energy absorbed by photosystem II that is used up in photosynthesis (Lazár, 2015). Under control conditions, no major differences were observed among Col and single mutant plants. However, the quantum yield of the *ahk2 ahk3* double mutant was reduced, when compared to wild type and *ahk2* cultivated under control conditions at 14 DODS (Fig. 5C), which points to the decreased ability of absorbed energy usage in linear electron transport in primary photosynthesis. There was no difference in QY between *ahk3* and *ahk2 ahk3* in control conditions (Fig. 5C). However, exposure to drought masked the differences among the used lines, and there were no differences in quantum yield among plant genetic lines at 14 DODS. We also examined the effect of drought stress on individual plant lines. When compared to plants grown under control conditions, exposure to drought stress reduced the quantum yield of wild type and *ahk* single mutants. With the *ahk2 ahk3* double mutant, there was no significant difference between plants grown under control or drought conditions (Fig. 5C). The quantum yield often decreases in response to abiotic stress and may act as a marker for evaluating the ability to resist abiotic conditions for various plant mutant lines (Yao *et al*., 2018) or crop varieties (Findurová *et al*., 2023).

Color change is also a key determinant, using which we can predict the genotype of the observed plant. The most important was hue R57 G71 B46 (Fig. 5A). This hue corresponds to a darker color. This points at changes in the presence of photosynthetic pigments. No differences were observed among wild type and single mutants under control conditions. The same result was detected with wild type and single mutants exposed to drought stress. In *ahk2 ahk3* double mutant the R57 G71 B46 value was strongly decreased under drought stress conditions, when compared to other plant lines (Fig. 5D). Under control conditions the R57 G71 B46 value for the *ahk2 ahk3* double mutant was decreased only when compared to wild type and the *ahk2* single mutant. Drought stress increased the R57 G71 B46 value in all observed plant lines. The presence of cytokinin has a strong effect on the chlorophyll content of leaves (Richmond and Lang, 1957) and treatment with different types of cytokinin modulated the chlorophyll *a*, chlorophyll *b* content as well as their ratio in cultivated apple leaves *in vitro* (Dobránszki and Mendler-Drienyovszki, 2014). A strong decrease in chlorophyll (to approximately 70% of wild type) was detected in the *ahk2 ahk3* double mutant (Riefler *et al*., 2006). However, studies linking color segmentation to the content of photosynthetic pigments are still absent.

In conclusion, our results indicate that *ahk2* and *ahk3* show only low resistance to stress in the early phases of drought. The results seem to be partially different from previously published drought tolerant phenotype of *ahk2* and *ahk3* (Tran *et al*., 2007; Kang *et al*., 2012) which can be caused by different experimental setup. Previously published research dealt with long-term stress combined with recovery whereas our work focused on early stages of the drought stress.

### Future Perspectives

Plant phenotyping with the help of state-of-the-art imaging techniques is a rapidly developing field that shows enormous potential in terms of the ability to recognize individual plant phenotypes that have improved resistance to both abiotic and biotic stresses. The new techniques being developed will provide more accurate information about how individual plants react to the applied stress and will further help us better understand a whole range of mechanisms that are responsible for the given reaction. Kinetic imaging of chlorophyll fluorescence is a very sensitive method that provides a whole range of information about plant physiology and fitness. There are also spectral analyses where, with the help of hyperspectral cameras, we can obtain information about the composition of individual pigments in plants or other metabolites. Raman spectroscopy then expands the spectrum of analysed substances by including more complex organic compounds. X-ray tomography makes it possible to investigate internal structures of the plant and monitor the morphology inside the observed objects. Further, 3D modelling with the help of laser scanners, increases the accuracy of the 3D model and thus the possibility of obtaining precise information about plant morphology. We can also expect that new non-invasive sensors with higher sensitivity and precision will soon become available and will provide more accurate information needed for plant phenotyping. Moreover, new developing AI approaches will fundamentally enhance individual analyses. These new measurement techniques and analytical methods will significantly improve our ability to uncover new plant phenotypes and traits associated with environmental stresses.

## Supplementary data

The following supplementary data are available at JXB online.

Fig. S1. The correlogram showing the data multicollinear nature.

Fig. S2. The supporting data for PCAs.

Fig. S3. The parameter contribution in PCA of global data.

Table S1. List and description of parameters obtained from FV/FM protocol.

Table S2. List and description of parameters obtained from Kautsky slow induction kinetics protocol. Table S3. List and description of parameters obtained from Quenching analysis protocol.

Table S4. List and description of parameters obtained from Rapid light curve protocol. Table S5. Confusion matrices for random forest for stress classification on total data. Table S6. Confusion matrices for random forest for stress classification on 5 DODS. Table S7. Confusion matrices for random forest for stress classification on 14 DODS. Table S8. Confusion matrices for random forest for genotype classification on 14 DODS. Dataset S1. Morphological, RGB and fluorescence data.

## Acknowledgements

We would like to acknowledge the Plant Sciences Core Facility and Biological Data Management and Analysis Core Facility of CEITEC Masaryk University (funded by ELIXIR CZ research infrastructure MEYS Grant No: LM2023055) for their valuable technical support.

## Author contributions

All authors conceptualized the project. JŠ performed the phenotyping experiments. JR analysed the data. PPS prepared figures. All authors evaluated the data, wrote the manuscript, contributed to reviewing the manuscript and approved the submitted version.

